# Molecular bases for USP48 cis-activity regulation and hyperactivation by Cushing’s disease-associated mutations

**DOI:** 10.64898/2025.12.27.696724

**Authors:** Gai Ando, Yasutaka Tsujimoto, Kei Moritsugu, Keijun Kakihara, Akinori Kidera, Shozo Yamada, Masayuki Komada, Hidenori Fukuoka, Toshiaki Fukushima

**Author notes:** equal contribution (Gai Ando, Yasutaka Tsujimoto). Toshiaki Fukushima and Kei Moritsugu; **Email:**. **Author Contributions:** K. M., A. K., H. F., and T.F. designed research; G. A., Y. T., and K. K. performed research; K. K. and T. F. contributed new reagents or analytic tools; S. Y. contributed clinical samples; G. A., Y. T., K. M., A. K., H. F., and T. F. analyzed data; K. M., H. F. and T. F. drafted the manuscript; M. K. and A. K. critically revised the manuscript. Competing Interest Statement: No competing interest.

## Abstract

Ubiquitin-specific protease 48 (USP48) is a deubiquitinating enzyme involved in various pathophysiological processes, including DNA repair, tumorigenesis, and inflammation. Somatic mutations in *USP48* (M415I/V) are associated with pituitary tumors in Cushing’s disease. These variants lead to increased USP48 enzyme activity and adrenocorticotropic hormone (ACTH) hypersecretion, a key feature of Cushing’s disease, indicating a direct link to the condition. Using biochemical experiments and in silico structural analyses, we identified Y414 in the catalytic USP domain as a critical residue for USP48 activity. In the wild-type USP domain, the Y414 side chain was oriented to ‘closed,’ which led to catalytic triad misalignment and weak ubiquitin recognition. We also found that the C-terminal region (CTR) of USP48 is a cis-activating region. It interacts with the USP domain and ubiquitin tail, thereby facilitating the alignment of the catalytic triad and enhancing ubiquitin recognition ability. Additionally, genomic analyses of pituitary tumors in Japanese patients with Cushing’s disease (n=46) confirmed that USP48 M415I/V variants occurred at a frequency of 10%, which is consistent with previous reports. Our structural model suggests that M415I/V sterically interferes with and fixes the closed Y414 side chain to ‘open,’ resulting in catalytic triad alignment and increased ubiquitin recognition ability. Furthermore, USP48 with M415I/V showed enhanced CTR-mediated activity, indicating that the open Y414 side chain and CTR can cause hyperactivation. This study provides a molecular basis for the novel cis-activation regulation of USP48 and its hyperactivation by Cushing’s disease-associated variants.

**Significance Statement:** The regulatory mechanisms of DUB activity are not yet fully understood. This study identifies the CTR of USP48 as an activity-promoting region, a mechanism also present in USP7 and potentially other closely related DUBs such as USP40 and USP47. Despite the disease association of some DUBs, developing drugs to inhibit specific DUB members or variants remains challenging. The substitutions of USP48 M415 embedded in the USP domain to Ile/Val are associated with Cushing’s disease, which lacks adequate targeted therapeutics. This study reveals that these variants alter the orientation of Y414, leading to hyperactivation. These findings provide a molecular basis for developing USP48^M415I/V^-specific inhibitors for Cushing’s disease therapy.

## Introduction

Ubiquitination is a post-translational modification that occurs in more than half of cellular proteins (1). It induces degradation of the target proteins or affects their properties, including complex formation, subcellular localization, and enzyme activity. Deubiquitinating enzymes (DUBs) antagonize ubiquitination. The human genome encodes approximately 100 genes for DUBs, approximately half of which belong to the ubiquitin-specific protease (USP) subfamily (2). Many DUBs play roles that others cannot compensate for. The underlying mechanisms include different tissue expression patterns, distinct subcellular localization, and specificity for substrate proteins and ubiquitin chains (3). Additionally, some DUBs have conditional activation mechanisms, including cis-regulation by self-regulatory domains and trans-regulation by accessory proteins (4). Several DUBs are associated with human diseases and have attracted attention as potential drug targets (4).

USP48, a member of the USP family, is expressed in various tissues (5) and performs multiple pathophysiological functions. USP48 regulates DNA repair (6, 7) and cell cycle progression (8) and can either promote (9–12) or suppress (13–15) cancer growth and metastasis. USP48 also functions as a proinflammatory factor (16–18), whereas it functions as an anti-inflammatory factor in different contexts (19, 20). Recent studies have reported that genetic variants of *USP48* are associated with Cushing’s disease (21, 22) and hearing loss (23). As described in detail below, genetic variants of *USP48* in Cushing’s disease increase DUB activity (22), whereas some variants linked to hearing loss reduce DUB activity (23).

Cushing’s disease is caused by a pituitary tumor that secretes adrenocorticotropic hormones (ACTH), leading to hypercortisolism. Symptoms include moon face, central obesity, and complications such as diabetes, hypertension, immunocompromise, osteoporosis, muscle atrophy, and depression (24). The molecular mechanisms underlying ACTH hypersecretion remain unclear and no therapeutic agents have been developed to specifically target this pathophysiology or alleviate autonomous ACTH hypersecretion. The first-line treatment is pituitary tumor resection; however, the disease is intractable and has a poor prognosis (standardized mortality ratio ∼3.0) (24) Genomic analysis of Cushing’s disease tumors revealed that somatic variants of *USP48* and *USP8* occur at frequencies of ∼10% and ∼50%, respectively (25). Curiously, these variants are often found in pituitary tumors that secrete ACTH but are very rare in other tumors and cancers (COSMIC database; https://cancer.sanger.ac.uk). The resulting changes in amino acid sequence increase DUB activity (22, 26). Although the intermediate pathways are not fully understood, a recent study using iPSC-derived pituitary organoids showed that these variants increased the population of Tpit-positive ACTH-producing cells, the expression of the ACTH precursor *proopiomelanocortin* gene, and ACTH secretion (27). This indicates a causal link between these variants and the disease.

We previously showed that USP8 possesses an activity-regulation mechanism that is absent in other USP family members. USP8 has an autoinhibitory domain called the WW-like domain, which interacts with and inhibits the catalytic USP domain (28). The interaction between the WW-like and USP domains is stabilized by the phosphorylation-dependent binding of USP8 to 14-3-3 (28–30). Furthermore, we elucidated the molecular basis of USP8 hyperactivation in Cushing’s disease-associated variants. Most variants cause amino acid substitutions or deletions in the 14-3-3-binding motif (14-3-3-BM), reducing its ability to bind to 14-3-3 (26). This relieves autoinhibition, leading to hyperactivation (28). For USP48, although the full length is enzymatically active, losing regions other than the USP domain dramatically reduces its activity (16, 23, 31, 32), implying a cis-regulatory mechanism. Cushing’s disease-associated variants in *USP48* cause the substitution of M415 in the USP domain with Ile or Val, leading to hyperactivation (22). However, the molecular mechanisms underlying the cis-regulation of USP48 activity and hyperactivation by the M415I/V variants remain unknown.

In this study, we addressed these issues. We elucidated the mechanism through which the C-terminal region (CTR) of USP48 functions as a cis-activating region. We also examined the frequency of *USP48* variants in Japanese patients with Cushing’s. Further analyses revealed that the M415I/V variants modified the structure of the USP domain, increasing enzyme activity in the presence or absence of a CTR-mediated regulatory mechanism.

## Results

### USP48 NTR, USP, and DUSP1 domains form an enzyme module

We analyzed the structure of the catalytic USP domain of wild-type USP48. Similar to other USPs, the predicted structure had a right-hand shape consisting of thumb, palm, and fingers subdomains (Supplementary Figure 1A). The ubiquitin-binding pocket, formed by the palm and fingers subdomains, captures the core of the ubiquitin molecules conjugated to substrate proteins. The catalytic groove, which is the boundary between the thumb and palm subdomains, interacts with the C-terminal tail of ubiquitin. This groove contains an active site composed of a catalytic triad that hydrolyzes the isopeptide bond between the ubiquitin C-terminus and substrate protein, resulting in deubiquitination.

USP48 has USP, three tandem DUSP (domain present in USP), and UBL (ubiquitin-like) domains. The N-terminal region (NTR; aa 1-88) and C-terminal region (CTR; aa 1010-1035) are both nonstructural. We found that NTR and DUSP1 were required for the expression of the USP domain in cells (Figure 1A and B). The predicted structure of the region aa 1-568, containing the NTR, USP, and DUSP1 domains (Supplementary Figure 1B), suggests a putative interaction between the NTR and DUSP1 domains. These results indicated their role in stabilizing the USP domain. Therefore, in this paper, the region aa 1-568 is referred to as the ‘enzyme module.’

**Figure 1.**
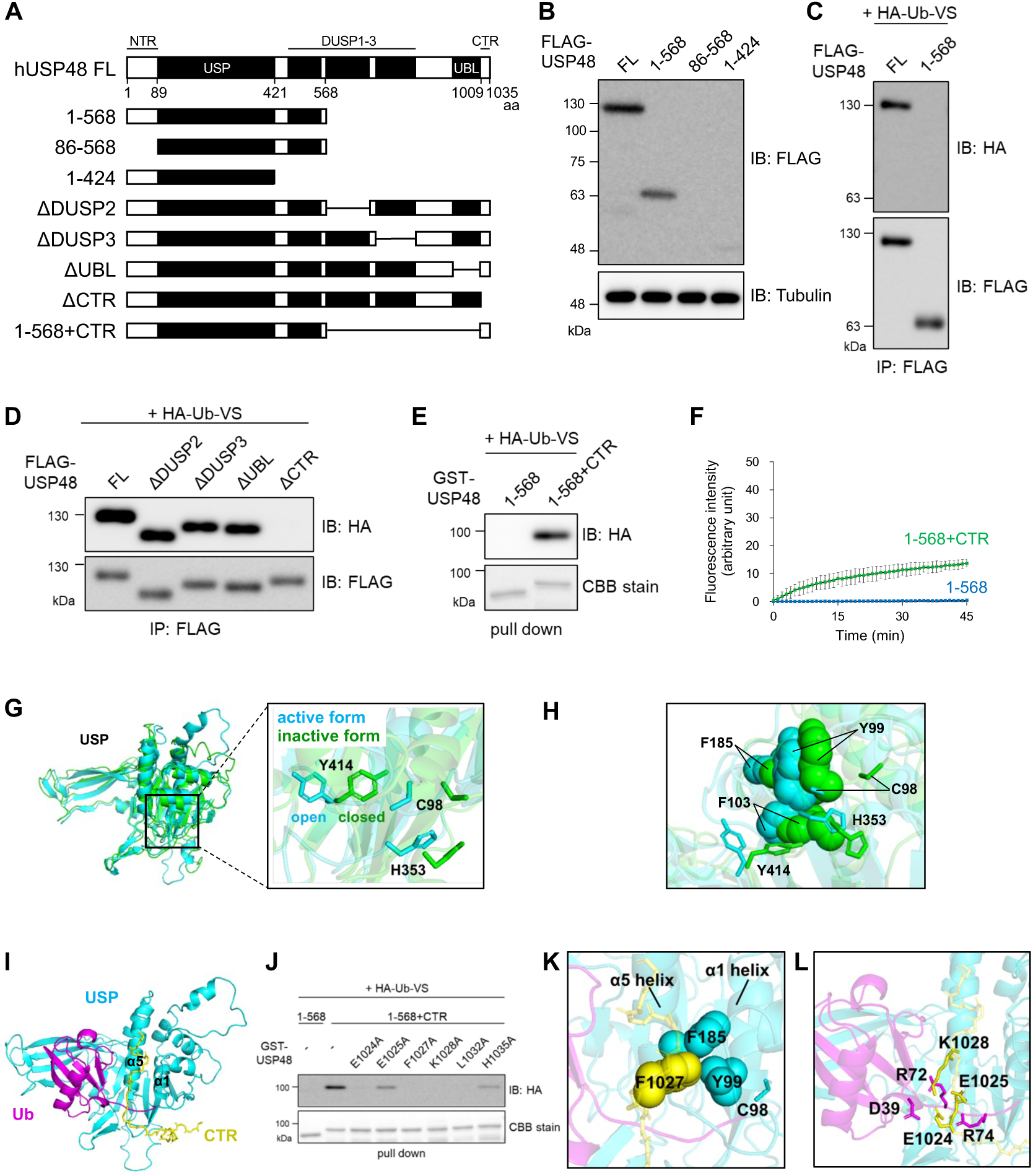
Auto-activation of USP48 via the CTR and the molecular basis. (A) Schematic structures of human USP48 and the truncated mutants used in B-F. NTR, N-terminal region; USP, USP domain; DUSP, domain present in ubiquitin-specific protease; UBL, ubiquitin-like domain; CTR, C-terminal region. (B) Expression levels of USP48 mutants. HEK293T cells expressing FLAG-tagged full-length USP48 (FL) or its mutants were lysed and subjected to immunoblotting (IB). (C-E) Profiling of USP48 mutants using an activity-based probe. In C-D, cells expressing FLAG-tagged USP48 mutants were lysed and subjected to immunoprecipitation (IP) with an anti-FLAG antibody. In E, GST-tagged USP48 mutants were expressed in E. coli, purified with glutathione-conjugated beads. The precipitates were labeled with HA-tagged-ubiquitin-vinyl sulfone (HA-Ub-VS) and subjected to IB and Coomassie Brilliant Blue (CBB) staining. (F) Deubiquitinating activity of USP48 mutants. GST-tagged USP48 mutants were subjected to an in vitro deubiquitination assay using ubiquitin 7-amido-4-methylcoumarin (Ub-AMC). The fluorescence intensity of the released AMC was measured. The graph shows the means ± standard deviations of three independent experiments. (G) Molecular dynamics (MD) simulations of the apo-structure of the USP domain of USP48 predicted by AlphaFold2 (AF2). The right panel is the enlarged view around the active groove. The initial structure (cyan) showed an active form, whereas the stable structure after MD simulation (green) showed an inactive form. The orientation of the Y414 side chain in the active and inactive forms is defined as ‘open’ and ‘closed,’ respectively. (H) Dislocation of hydrophobic residues near the active site. (I) The complex model comprising the USP domain (cyan), CTR (yellow), and ubiquitin (magenta) predicted by AF2. (J) Validation of the model by activity-based probe profiling. USP48 harboring mutations in the CTR were expressed in E. coli, purified with glutathione-conjugated beads, labeled with HA-Ub-VS, and subjected to IB and CBB staining. (K) Molecular structure around the active groove in the complex model. (L) Binding sites of CTR and ubiquitin in the complex model.

### The USP48 enzyme module is inactive by itself and activated by the CTR

Ubiquitin vinyl sulphone (Ub-VS) is an activity-based probe that covalently binds to the active site of the cysteine protease DUBs via the VS group. The probe labeled full-length USP48 but not the enzyme module (Figure 1C). The DUSP2-3 or UBL domains did not affect probe labeling; however, CTR was required (Figure 1D). Linkage of CTR to the enzyme module restored probe labeling (Figure 1E). The DUB activity assay using ubiquitin 7-amido-4-methylcoumarin (Ub-AMC) as a substrate also showed that the linkage of CTR to the enzyme module increased the enzyme activity (Figure 1F). These data demonstrate that the USP48 enzyme module is inactive on its own, but is activated by CTR.

### The molecular basis underlying that the USP48 enzyme module alone is inactive

We predicted the apo-structure of the USP domain using alpha-fold 2 (AF2) (33) and analyzed it using molecular dynamics (MD) simulations (Figure 1G). Probably because the prediction was influenced by the active structures of the other USPs, the initial structure had an active form with an aligned catalytic triad, including C98 and H353. However, during the simulations, it quickly transitioned to an inactive form, in which the catalytic triad was not aligned. The frequency of the catalytic triad alignment (P_aligned_) was 0.13 (Table 1, USP^WT^). In addition, we found that the conformation of the Y414 side-chain χ1 was simultaneously converted from trans to gauche+ (Figure 1G, right panel, Supplementary Figure 2) and defined its orientation in the active and inactive forms as ‘open’ and ‘closed,’ respectively. The frequency of Y414 side-chain opening (P_open_) was 0.0 (Table 1, USP^WT^).

**Table 1.** The frequency of Y414 side-chain opening, the frequency of the catalytic triad alignment, and the root mean square deviation (RMSD) of ubiquitin from the predicted USP-Ub structure were calculated from MD simulations for the indicated structures.

| Structure | Mutation in USP | Frequency of Y414 side-chain opening ($P_{open}$) | Frequency of catalytic triad alignment ($P_{aligned}$) | RMSD of Ub (Å) |
| --- | --- | --- | --- | --- |
| USP | WT | 0.0 | 0.13 | - |
|  | Y414A | - | 0.38 | - |
|  | M415I | 1.0 | 0.82 | - |
| USP-Ub | WT | 0.038 | 0.058 | $3.6 \pm 0.46$ |
| | Y414A | - | 0.51 | $3.6 \pm 0.42$ |
| | M415I | 0.89 | 0.95 | $2.7 \pm 0.50$ |
| USP-CTR-Ub | WT | 0.21 | 1.0 | $2.7 \pm 0.41$ |
| | Y414A | - | 1.0 | $2.9 \pm 0.52$ |
| | M415I | 0.90 | 0.90 | $1.8 \pm 0.53$ |

Y414 is located on the β-sheet of the palm subdomain, which is distant from the catalytic triad (Supplementary Figure. 1A). The causal relationship between the closed Y414 side chain and the catalytic triad misalignment is shown in Figure 1G and H. The transition to the closed Y414 side chain leads to a shift in the surrounding hydrophobic residues (Y99, F103, and F185 on the α5 helix) and the subsequent dislocation of the catalytic residues (C98 and H353) on the active site.

The MD simulations showed a small P_aligned_ for both the wild-type and Y414A mutation, indicating the requirement of an open Y414 side chain for the catalytic triad alignment (Table 1, USP^Y414A^). Together, these data indicate that the USP48 enzyme module alone is inactive because the Y414 side chain is preferentially oriented to close, leading to catalytic triad misalignment.

### The CTR facilitates catalytic triad alignment and ubiquitin recognition by the USP domain

We constructed a USP-CTR-Ub ternary complex model by use of AF2 (Figure 1I). CTR binds to the USP domain and ubiquitin. We investigated the effect of mutations to Ala in the predicted binding site on the CTR (<4 Å interatomic distance). These mutations reduced Ub-VS labeling of the CTR-linked enzyme module (Figure 1J), confirming the validity of the model.

A closer look at the model revealed that F1027 in the CTR interacts with multiple hydrophobic amino acid residues in the active groove of the USP domain (Figure 1K), possibly contributing to the allosteric catalytic triad alignment. In addition, ubiquitin recognition seems to be facilitated by multiple residues in CTR (E1024, E1025, and K1028) (Figure 1L). E1024 forms a salt bridge with R72 in the ubiquitin tail. E1025 interacts with the main chain of R74. K1028 forms a salt bridge with D39 in the ubiquitin core, which complements the shallow ubiquitin-binding pocket.

To investigate the effect of CTR on the USP domain structure, we compared models of the USP-Ub binary complex and the USP-CTR-Ub ternary complex. MD simulations revealed that CTR slightly increased P_open_ and significantly increased P_aligned_. Furthermore, calculation of the root mean square deviation (RMSD) of ubiquitin from the predicted USP-Ub structure revealed that CTR also assists in capturing ubiquitin (Table 1, USP^WT^-Ub and USP^WT^-CTR-Ub). It was supported by the calculated binding free energies ΔGbind between the USP domain and ubiquitin (USP-Ub, -72.5± 9.9 kcal/mol; USP-CTR-Ub, -102 ± 16 kcal/mol). These results indicated that CTR facilitates catalytic triad alignment and ubiquitin recognition.

### Japanese patients with Cushing’s disease have USP48 M415I/V variants as frequently as previously reported

We performed genomic analysis of pituitary tumors resected from Japanese patients with Cushing’s disease by Sanger sequencing, targeting *USP48* exon 10, which encodes the USP domain. Among the 100 patients, genetic variants were found in 8 patients (8%). These pathogenic variants cause M415I/V substitutions in the USP48 USP domain, similar to previously reported mutation patterns (25). The clinical characteristics of the patients are summarized in Table 2. *USP48* variants were predominantly observed in females (73.9%). Compared to the group with wild-type *USP48*, these patients exhibited lower ACTH levels in both early morning and late night measurements. Additionally, they showed lower ACTH levels in the low-dose dexamethasone suppression test (LDDST) and smaller tumors.

**Table 2.** The clinical characteristics of the patients with Cushing’s disease.

|  | patients with<br>USP48_M415I/V<br>(n=5) | (n) | patients with<br>wild-type USP48<br>(n=41) | (n) | p value |
| --- | --- | --- | --- | --- | --- |
| sex (female, %) | 100.0 | 5 | 65.9 | 27 | 0.30 |
| age (years) | 48.0 ± 18.3 | 5 | 46.4 ± 17.6 | 41 | 0.85 |
| BMI (kg/m2) | 26.4 ± 3.5 | 4 | 26.3 ± 4.2 | 26 | 0.95 |
| early morning cortisol (µg/dL) | 22.3 ± 6.3 | 5 | 19.8 ± 9.2 | 41 | 0.57 |
| early morning ACTH (pg/mL) | 41.9 ± 20.3 | 5 | 104.6 ± 70.4 | 41 | 0.06 |
| late night cortisol (µg/dL) | 14.4 ± 8.0 | 5 | 17.0 ± 7.6 | 41 | 0.47 |
| late night ACTH (pg/mL) | 24.4 ± 5.9 | 4 | 101.9 ± 76.2 | 41 | 0.05 |
| UFC (µg/day) | 608.8 ± 706.0 | 5 | 492.3 ± 673.5 | 33 | 0.72 |
| LDDST cortisol (µg/dL) | 16.3 ± 7.7 | 5 | 18.9 ± 12.6 | 38 | 0.66 |
| LDDST ACTH (pg/mL) | 37.7 ± 34.9 | 4 | 86.6 ± 81.4 | 33 | 0.25 |
| tumor size (mm) | 6.6 ± 2.6 | 5 | 14.2 ± 13.4 | 35 | 0.22 |

Available data from the indicated number of cases are shown for each parameter.
Abbreviation: BMI, body mass index; ACTH, adrenocorticotrophic hormone; UFC, urinary free cortisol; LDDST, low-dose dexamethasone suppression test.

### The enzyme module of USP48^M415I/V^ has enzyme activity by itself and is further activated by the CTR

M415 is located on the β sheet of the palm subdomain, next to Y414 (Supplementary Figure 1A). USP48^M415I/V^ was more strongly labeled with Ub-VS compared to wild-type USP48 (Figure 2A), indicating high enzyme activity, which is consistent with previous reports (22). USP48^M415I^ expressed in E. coli was also strongly labeled with Ub-VS (Figure 2B), suggesting that hyperactivation of USP48^M415I/V^ may occur without other binding proteins. Unlike the wild-type enzyme module, the USP48^M415I^ enzyme module was labeled with Ub-VS (Figure 2C) and showed enzymatic activity (Figure 2D). CTR enhanced probe labeling of USP48^M415I^ (Figure 2E and F) and its enzyme activity (Figure 2G). These data indicate that the enzyme module of USP48^M415I/V^ has enzyme activity and is further activated by CTR.

**Figure 2.**
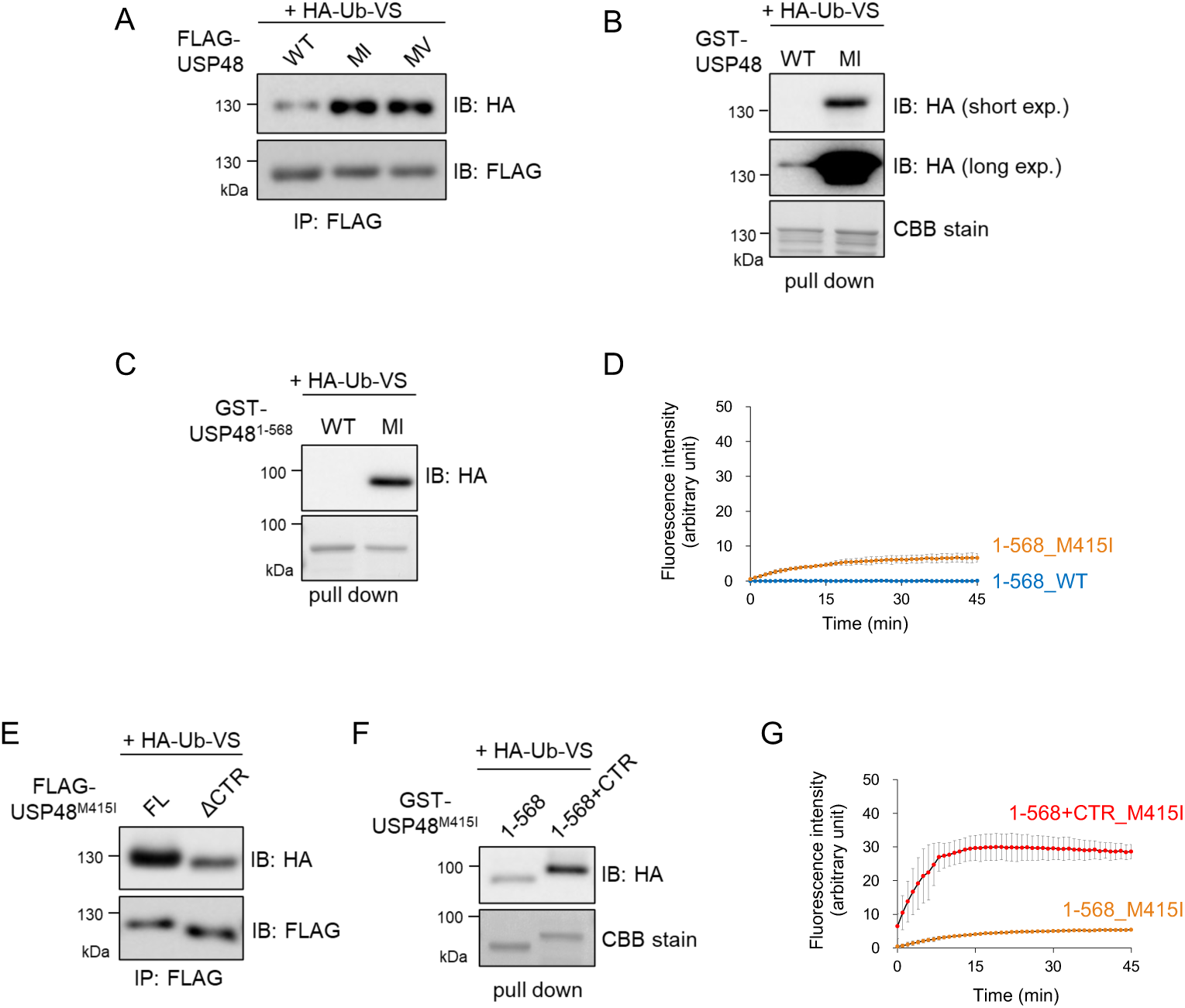
Hyperactivation of USP48 by Cushing’s disease-associated mutations. (A-C) Effects of M415I/V mutations on the activity-based probe profiling of USP48. In A, cells expressing USP48 or USP48^M415I/V^ were lysed and subjected to IP with an anti-FLAG antibody. In B-C, USP48 (FL or 1-568) or M415I mutants were expressed in E. coli, precipitated with glutathione-conjugated beads. The precipitates were labeled with HA-Ub-VS and subjected to IB and CBB staining. (D) Effects of M415I mutation on USP48^1-568^ enzyme activity. USP48^1-568^ or its M415I mutant were expressed in E. coli, purified with glutathione beads, and subjected to Ub-AMC assay. The graph shows the means ± standard deviations of three independent experiments. (E-F) Influence of the CTR on the activity-based probe profiling of USP48^M415I^. Experiments similar to those in A-C were performed. (G) Influence of the CTR on USP48^1-568,M415I^ enzyme activity. Similar experiments to those in D were performed.

### M415I/V variants interfere with the closed Y414 side chain and change the orientation to open, which allows the catalytic triad alignment

We compared the effects of the M415I/V/L mutations. Notably, the M415L mutation has not been reported in Cushing’s. M415I/V mutations increased the activity of the CTR-linked enzyme module (Figure 3A), whereas the M415L mutation had no effect (Figure 3B). We placed Ile/Val/Leu at M415 of the structural model with a closed Y414 side chain. The β-branched amino acid Ile/Val appeared to interfere with the closed Y414 side chain sterically, whereas the γ-branched amino acid Leu did not (Figure 3C). These data suggest that when M415I/V mutations occur, the closed Y414 side chain sterically interferes and opens.

**Figure 3.**
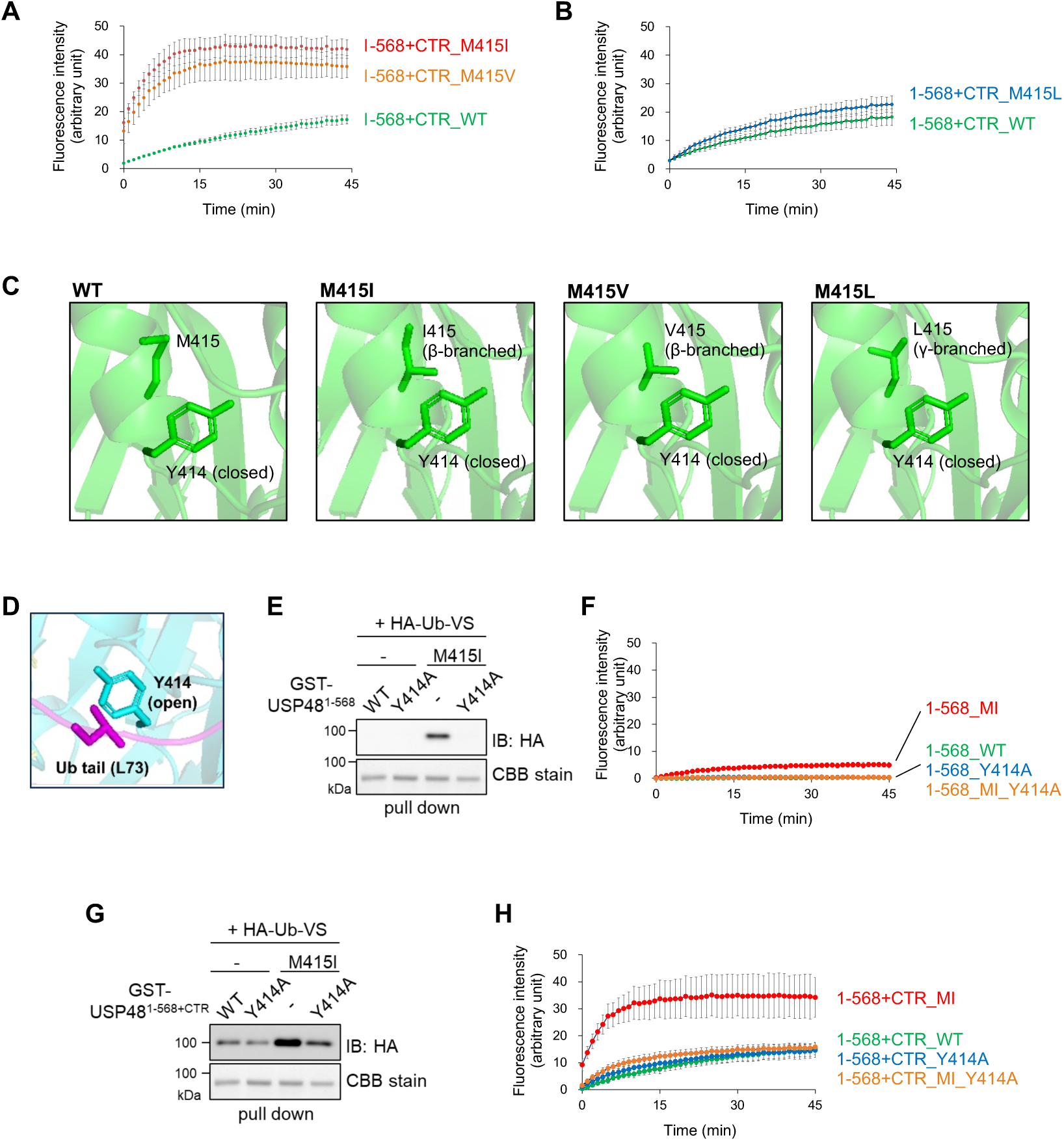
Molecular basis of the hyperactivation of USP48 by Cushing’s disease-associated mutations. (A-B) Effects of substitution of M415 with Cβ- or Cγ-branched amino acids on enzyme activity. The indicated proteins were expressed in E. coli, purified using glutathione beads, and subjected to the Ub-AMC assay. The graph shows the means ± standard deviation of three independent experiments. (C) Interference of Ile or Val placed at M415 with a closed Y414 side chain. The images show the structural model after MD simulation with the closed Y414 side chain and those in which M415 was replaced with Ile, Val, and Leu. (D) Ubiquitin recognition is facilitated by the open Y414 side chain of USP48^M415I^. A complex model comprising the USP domain of USP48^M415I^ (cyan) and ubiquitin (magenta) was predicted using AF2, which indicated the proximity of the open Y414 side chain to ubiquitin. (E-H) Effects of the Y414A mutation on activity-based probe profiling and enzyme activity of USP48. In E and G, USP48 (1-568 or 1-568+CTR) or Y414A mutants were expressed in E. coli and precipitated using glutathione-conjugated beads. The precipitates were labeled with HA-Ub-VS and subjected to IB and CBB staining. In F and H, the precipitates were subjected to the Ub-AMC assay. The graph shows the means ± standard deviation of three independent experiments.

**Figure 4.**
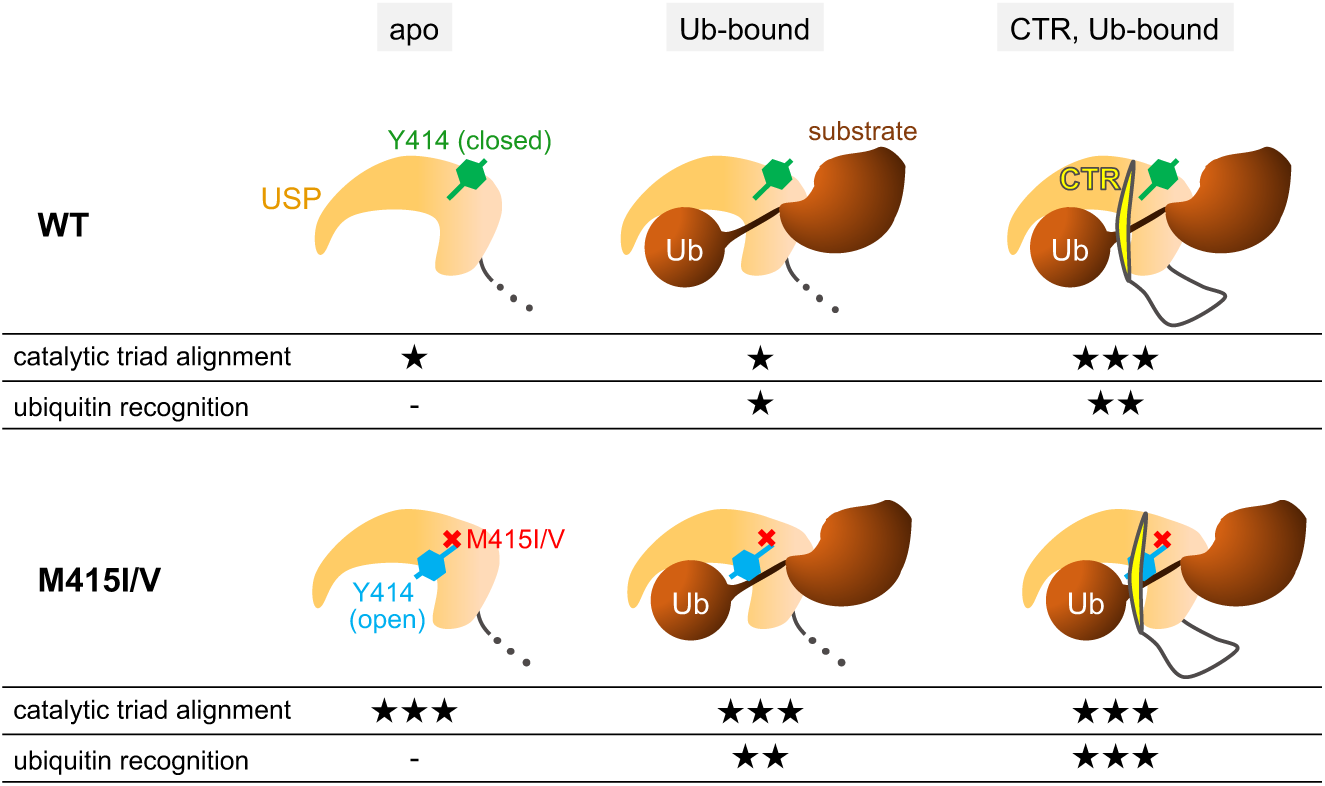
Molecular mechanisms of USP48 activity regulation and its hyperactivation by Cushing’s disease-associated mutations (working hypothesis). See text for details. The number of * is based on Table 1.

We investigated the effect of the M415I mutation on the USP domain structure using MD simulations (Table 1, USP^M415I^). Consistent with the above data, the M415I mutation increased P_open_ to 1.0 and P_aligmed_ to 0.82, indicating that USP48^M415I^ has an open Y414 side chain and an aligned catalytic triad without ubiquitin or CTR.

### The open Y414 side chain contributes not only to the catalytic triad alignment but also to the ubiquitin recognition

In a USP-Ub binary complex model, the open Y414 side chain was in contact with the ubiquitin tail (L73, interatomic distance < 4 Å) (Figure 3D). The Y414A mutation in the wild-type enzyme module did not affect Ub-VS labeling or enzyme activity, whereas the Y414A mutation in the M415I-type enzyme module reduced Ub-VS labeling and enzyme activity (Figure 3E and F). Similarly, when we used the CTR-linked enzyme modules, the Y414A mutation in the wild-type had no effect, whereas the Y414A mutation in the M415I-type reduced Ub-VS labeling and enzyme activity (Figure 3G and H). MD simulations showed that the RMSD of ubiquitin from the bound structure with the Y414A mutation was larger than that with the M415I mutation, irrespective of whether the CTR was unbound (Table 1, USP^Y414A^-Ub, USP^M415I^-Ub) or bound (Table 1, USP^Y414A^-CTR-Ub, USP^M415I^-CTR-Ub) to the USP. Therefore, the reduced enzyme activity in the M415I-type enzyme module caused by the Y414A mutation was due to weak ubiquitin recognition ability.

MD simulations of the USP-Ub binary complex model showed that the M415I mutation not only increased P_aligned_ but also decreased the RMSD of ubiquitin (Table 1, USP-Ub, USP^M415I^-Ub), indicating a higher ubiquitin recognition ability. MD simulations of the USP-CTR-Ub ternary complex model also showed that the M415I mutation decreased the RMSD of ubiquitin (Table 1; USP-CTR-Ub, USP^M415I^-CTR-Ub). These data indicate that the open Y414 side chain in USP48^M415I^ contributes not only to catalytic triad alignment, but also to ubiquitin recognition.

## Discussion

Our study revealed that wild-type USP48 has a closed Y414 side chain, leading to misalignment of the catalytic triad and weak ubiquitin recognition. We also identified the CTR of USP48 as an activity-promoting region. One of the hearing loss-associated USP48 variants lacking aa1020-1035 showed no enzymatic activity (23). Our study suggests that this is due to CTR dysfunction. In our USP-CTR-Ub ternary complex model, the USP domain and CTR sandwiched the ubiquitin tail. Therefore, during deubiquitination, it is likely that the USP domain-ubiquitin complex is formed first, followed by the addition of CTR to form a ternary complex. Our data indicate that adding CTR increases the catalytic triad alignment and ubiquitin recognition ability, leading to the progression of the deubiquitination reaction.

Molecular evolutionary analysis has shown that the USP domain of USP48 is closely related to those of USP7, USP40, and USP47 (34). Intriguingly, the regulation of USP48 activity resembled that of USP7, a well-known DUB of the E3 ligase Mdm2, which targets p53. The USP domain of USP7 is inactive on its own. This is because the catalytic triad is not properly aligned due to the electrostatic network in the switching loop and packing by hydrophobic residues around the active groove (35). Similarly to USP48, USP7 is activated by its CTR: the CTR binds to the α5 helix and switching loop in the USP domain, filling in the loose hydrophobic packing and aligning the catalytic triad (36). There was no homology between the amino acid sequences of USP7 and USP48 CTRs, indicating that they were optimized to fit their respective USP domains. It would be interesting to investigate whether other closely related DUBs (USP40 and USP47) have similar activity regulation mechanisms.

Some DUBs are known in which cis-regulatory mechanisms are modulated by other factors, such as post-translational modification, interaction with trans-regulatory proteins, inclusion in particular protein complexes, and localization to specific organelles (4, 28, 37, 38). We speculated that the CTR of USP48 might more readily bind to the USP domain in the presence of certain factors, leading to a remarkable increase in enzyme activity. In response to cytokine stimulation, USP48 is activated via phosphorylation and regulates inflammatory signaling by deubiquitinating TRAF2 and NF-κB p65 (16, 18). Upon DNA damage, USP48 is recruited to the damage site and reduces the multi-mono-ubiquitination of histone H2A, thereby regulating DNA repair. USP48 is activated by interactions with non-target ubiquitin molecules of H2A (7). Thus, it would be interesting to examine whether phosphorylation and binding to auxiliary ubiquitin facilitate CTR interaction with the USP domain.

Genomic analyses of pituitary tumors in Japanese patients with Cushing’s disease have identified USP48 M415I/V variants. The frequencies of these variants, particularly their higher occurrence in females, are consistent with previously reported data (25). While one study found no significant differences in the clinical phenotypes between patients with Cushing’s disease with and without USP48 variants (21), other studies indicated that patients with USP48 variants had smaller tumor sizes (22, 39). Our cohort revealed that patients with USP48 variants presented with smaller tumor sizes and lower ACTH levels in the early morning, late night, and LDDST. These findings suggest that patients with USP48 variants may exhibit mild clinical features. Further research is necessary to understand why patients with the USP48 variant have these characteristics.

In this study, we elucidated the mechanism of hyperactivation of USP48^M415I/V^. M415I/V, embedded in the USP domain, sterically interferes with Y414 and keeps it open, resulting in catalytic triad alignment and increased ubiquitin recognition ability. Similar to wild-type USP48, USP48^M415I/V^ can undergo CTR-mediated enhancement of activity. Given that USP-Ub binary complex formation may be a prerequisite for CTR interaction, CTR-mediated activity enhancement is more likely to occur in USP48^M415I/V^ because of their high ubiquitin recognition ability. Finally, both the open Y414 side chain and CTR cause hyperactivation. This study provides a molecular basis for the developing of USP48^M415I/V^-specific inhibitors for Cushing’s disease therapy.

The intermediate pathways through which mutations in USP48 or USP8 cause ACTH hypersecretion are not fully understood. Previous reports have suggested the involvement of EGFR, Hedgehog, and NF-κB pathways (21, 22, 26, 27). However, because the same genetic variants of *USP48* or *USP8* as those in Cushing’s disease have rarely been observed in other tumors and cancers, corticotroph-specific proteins/pathways other than these ubiquitous pathways may also be involved. Although the physiological substrates of USP48 and USP8 appear entirely different, their hyperactivation by mutations is associated with Cushing’s disease. In this study, we found that wild-type USP48 activity was switched on and off by a CTR-mediated mechanism, whereas USP48^M415I/V^ exhibited enzymatic activity to some extent without a CTR-mediated mechanism. Abnormalities in the conditional activation mechanism may allow USP48^M415I/V^ to deubiquitinate proteins that differ from their physiological substrates, raising the possibility that Cushing’s disease-associated USP48/USP8 mutants may have common disease-specific substrates. A search for substrates is required to understand the pathogenic mechanisms of Cushing’s disease.

## Materials and Methods

### Plasmid construction

Human USP48 cDNA was amplified by PCR from HeLa cells. The N-terminal FLAG-tagged USP48 plasmid was constructed by subcloning USP48 cDNA into the pFLAG-CMV2 vector (Sigma-Aldrich). The N-terminal GST-tagged USP48 plasmid was constructed by subcloning into a pGEX-6P1 vector (GE Healthcare). Mutagenesis was achieved using a Prime STAR Mutagenesis Basal Kit (Takara).

### Protein preparation

For Ub-VME labeling and Ub-AMC assay, GST-tagged USP48 (wild-type and mutants) were produced in Escherichia coli Rosetta (DE3) pLysS cells cultured overnight at 18 °C. Cells were lysed in ice-cold lysis buffer (50 mM Tris-HCl, pH 7.4; 150 mM NaCl; 50 mM NaF and 1% Triton X-100) supplemented with protease inhibitors (1 μg/ml each of aprotinin, leupeptin, and pepstatin A). After sonication and centrifugation, the supernatants were incubated with glutathione sepharose 4B (GE Healthcare). Post 1 h incubation at 4 °C, beads were washed thrice with lysis buffer and subsequently incubated in ice-cold elution buffer A (50 mM Tris-HCl, pH 8.0; 1 mM EDTA; 1 mM dithiothreitol (DTT) and 10 mM reduced glutathione; Wako) for 30 min at 4 °C. The purity and concentration of eluates were validated using SDS-PAGE and Coomassie Brilliant Blue (CBB) staining, with Quick CBB (Wako). FLAG-tagged USP48 (wild-type and mutants) were expressed in HEK293T cells. Cell lysis and immunoprecipitation were performed as described below. USP48 levels in each sample were compared using CBB staining and immunoblotting, with an equal number of USP48 molecules used in the assay.

### Cell culture and transfection

HEK293T cells were grown in high-glucose Dulbecco’s modified Eagle’s medium (Nacalai Tesque) supplemented with 10% fetal bovine serum (FBS), 100 units/ml of penicillin, and 0.1 mg/ml streptomycin at 37 °C and 5% CO_2_. Plasmid transfection was performed using polyethyleneimine (Polyscience) following standard protocol. The cells were lysed 48 h after transfection.

### Immunoprecipitation and immunoblotting

Cells were lysed in ice-cold lysis buffer supplemented with protein inhibitors, and the supernatants were collected after centrifugation. For immunoprecipitation, anti-FLAG M2 antibody-conjugated agarose beads (Sigma-Aldrich) were used following the manufacturer’s instructions. After immunoprecipitation, the beads were washed thrice with lysis buffer. For the elution of FLAG-tagged proteins, the beads were incubated in TBS with 200 ng/μl FLAG peptide (Sigma Aldrich) at 4 °C for 30 min.

For immunoblotting, samples were incubated in SDS-PAGE sample buffer (62.5 mM Tris-HCl, pH 6.8; 2% SDS; 5% 2-mercapto ethanol; 10% glycerol; and 0.1 mg/ml bromophenol blue) at 98 °C for 5 min or at 37 °C for 30 min. The samples were then separated using SDS-PAGE. Immunoblotting was performed according to the standard procedures. The primary antibodies used for immunoblotting were as follows: anti-FLAG (clone M2, Sigma Aldrich), anti-HA (clone 3F10, Roche), and anti-α-tubulin (clone 10G10, Wako) antibodies. The secondary antibodies used were peroxidase-conjugated anti-mouse, anti-rat, and anti-rabbit IgG antibodies (GE Healthcare). Proteins were detected using ECL Prime Western Blotting Detection Reagents (GE Healthcare) and ImageQuant LAS 4000 Mini (GE Healthcare).

### HA-Ub-VS labeling

GST/FLAG-tagged USP48 (wild type and mutant) was incubated with 1 μM HA-tagged Ub-VS (HA-Ub-VS; R&D Systems) in TBS supplemented with 1 mM DTT at 37 °C for 30 min. The samples were subjected to immunoblotting.

### Ub-AMC assay

GST-tagged USP48 (wild-type and mutants) were mixed with 1 μM Ub-AMC (Peptide Institute) and 10 mM DTT in TBS in 96-well medium-binding, flat-bottom, black plates (Thermo Scientific). Plates were set on Varioskan LUX (Thermo Scientific) and incubated at 37 °C. During incubation, the fluorescence intensity of AMC released from ubiquitin was measured at 1 min intervals using excitation and emission wavelengths of 345 and 445 nm, respectively. The fluorescence intensity of the sample without USP48 was measured as background and subtracted from each value.

### Structural modeling and MD simulation

The structural model of the USP48 catalytic domain (USP: aa 89-421) was obtained from the rank1 predicted model by AF2 (33). The sequence of the corresponding residues was used as the input, and standard parameters were used on the Colab Fold server (https://colab.research.google.com/github/sokrypton/ColabFold/). Similarly, two structural models of USP complexed with ubiquitin (Ub: aa1-76, the binary complex was named herein as USP-Ub) and USP-Ub complexed with the CTR (the ternary named herein as USP-CTR-Ub) were also constructed. Additional structures of the two mutants, Y414A and M415I, were modeled for USP, USP-Ub, and USP-CTR-Ub. Nine systems were used for the subsequent MD simulations and derived MD trajectory analyses.

For each system, a rectangular simulation box was constructed with a 12 Å margin to the box boundary and filled with TIP3P water molecules (40), along with sodium ions and chloride ions to achieve a concentration of ∼0.15 nM. The AMBER ff14SB was applied to the potential energy of the all-atom proteins (41, 42). MD simulations were performed using AMBER 20 (43) under constant temperature and pressure (NPT) conditions at *T* = 300 K and *P* = 1 atm using Berendsen’s thermostat and barostat (44) at a relaxation time of 1 ps. Electrostatic interactions were calculated using the particle mesh Ewald method (45). The simulation length was 1 μs, with a 2-fs time step, and bonds involving hydrogen atoms were constrained via the SHAKE algorithm (46). The coordinates were saved every 10 ps and used for analysis.

The frequency of Y414 side-chain opening, *P*_open_, was defined as the probability, through the last 800-ns MD simulation trajectory, that the side-chain torsion angle χ_1_ along N-Cα-Cβ-Cγ is in trans form (|χ_1_| > 150°). The frequency of catalytic triad alignment, *P*_align_, was defined as the probability, through the MD simulation trajectory, that the distance relating to two catalytic residues between the Cβ atom on C98 and the Cγ atom on H353 is less than 6 Å. This analysis did not include a third catalytic residue (N370) because a recent study argued that D371, rather than N370, functions as a catalytic residue (47). The RMSD of ubiquitin from the predicted USP-Ub structure after superimposing the USP palm subdomain (named herein as “RMSD of ubiquitin”) was calculated to evaluate the stability of Ub binding, because the predicted USP-Ub and USP-CTR-Ub structures were found to be the active Ub-binding forms by examining other USP crystal structures. The binding free energy of Ub was calculated using the MM-GB/SA module of AMBER 20 (43).

### DNA extraction and gene sequencing from pituitary tumors with Cushing’s disease

Genomic DNA was extracted from surgically resected pituitary tumor tissues using a QIAamp DNA FFPE Kit (QIAGEN). The coding region in exon 10 of the *USP48* gene was amplified by PCR using GoTaq DNA Polymerase (Promega) with the forward primer 5′-TGCCTGCTATAATCCTGGAAA-3′ and the reverse primer 5′-TCAGCAGAACCTTCTAAGTCTCA-3.’ The PCR products were purified using agarose gel electrophoresis and the QIAquick Gel Extraction Kit (QIAGEN), and sequenced using the BigDye Terminator Cycle Sequencing Kit on an ABI PRISM 310 Genetic Analyzer (Applied Biosystems).

### The clinical characteristics of the patients with Cushing’s disease

In all patients, Cushing’s disease was diagnosed based on the established guidelines (48) and confirmed using pathological findings. The clinical characteristics of the patients were retrospectively collected. Blood samples were obtained in the early morning, late night, and during LDDST, following established guidelines (48). The LDDST was performed using dexamethasone (1 or 0.5 mg) (48, 49). Plasma ACTH and serum cortisol levels were measured using enzyme immunoassays (TOSOH, RRID: AB_2783633 and AB_3076600). Urinary free cortisol (UFC) levels were determined using an immunoradiometric assay (IRMA) (TFB, RRID: AB_2894408) and chemiluminescent immunoassay (CLIA) (Siemens, RRID: AB_2893154). To ensure consistency, UFC levels measured by CLIA were adjusted to IRMA-equivalent values using the formula: Y = 0.917X − 2.80, where Y represents the IRMA value and X represents the CLIA value. The reference ranges are as follows: plasma ACTH, 7.2 to 63.3 pg/mL; serum cortisol, 7.07 to 19.6 μg/dL; UFC, 11.2 to 80.3 μg/day. Pituitary tumor size was measured by their diameters using magnetic resonance imaging (MRI). These methods were approved by the Research Ethics Committee of Kobe University Hospital (IRB No. 1363) and Moriyama Memorial Hospital (IRB No. 20006).

### Statistical analysis

Data are presented as means ± standard deviation. Categorical data were analyzed using Fisher’s exact test, whereas continuous data were evaluated using an unpaired t-test. Statistical analyses were performed using R version 4.1.3, with the significance level set at *p* < 0.05.

## Acknowledgments

We thank Drs. Naonobu Fujita, Hiroshi Iwasaki, and Hitoshi Nakatogawa at the Institute of Science Tokyo for their helpful discussions. We also acknowledge the help of the Biomaterials Analysis Division at the Institute of Science Tokyo for the DNA sequencing analysis. This work was supported by JSPS KAKENHI Grant Numbers JP19H05289 and JP21H00276 and AMED Grant Numbers JP22ym0126806j0001 and JP23ym0126806j0002 to T.F.; JSPS KAKENHI Grant Number JP21H02474 to M.K.; and Research Support Project for Life Science and Drug Discovery (BINDS) from AMED under Grant Number JP24ama121023 to K.M. and A.K.

## Notes

### Competing Interest Statement

The authors have declared no competing interest.

