## Supplementary Information for "Molecular bases for USP48 cis-activity regulation and hyperactivation by Cushing’s disease-associated mutations"

a. School of Life Science and Technology, Institute of Science Tokyo, 4259 Nagatsuta, Midori-ku, Yokohama, 226-8501, Japan.
b. Division of Diabetes and Endocrinology, Department of Internal Medicine, Kobe University Hospital, Kobe, 650-0017, Japan.
c. Graduate School of Science, Osaka Metropolitan University, 1-2 Gakuencho, Naka-ku, Sakai, Osaka, 599-8570, Japan.
d. Graduate School of Medical Life Science, Yokohama City University, 1-7-29 Suehirocho, Tsurumi, Yokohama, Kanagawa, 230-0045, Japan.
e. Cell Biology Center, Institute of Integrated Research, Institute of Science Tokyo, 4259 Nagatsuta, Midori-ku, Yokohama, 226-8501, Japan.
f. Institute of Molecular Biology, Academia Sinica, 128 Academia Road, Section 2, Nangang, Taipei, Taiwan.
g. Hypothalamic and Pituitary Center, Moriyama Memorial Hospital, Tokyo, 134-0081, Japan
1. equal contribution

*Toshiaki Fukushima and Kei Moritsugu

**Author Contributions:** K. M., A. K., H. F., and T.F. designed research; G. A., Y. T., and K. K. performed research; K. K. and T. F. contributed new reagents or analytic tools; S. Y. contributed clinical samples; G. A., Y. T., K. M., A. K., H. F., and T. F. analyzed data; K. M., H. F. and T. F. drafted the manuscript; M. K. and A. K. critically revised the manuscript.

**Competing Interest Statement:** No competing interest

**Keywords:** Deubiquitinating enzyme, ubiquitin-specific protease, ubiquitin, pituitary neuroendocrine tumor, Cushing's disease

**Supplementary Figure 1**

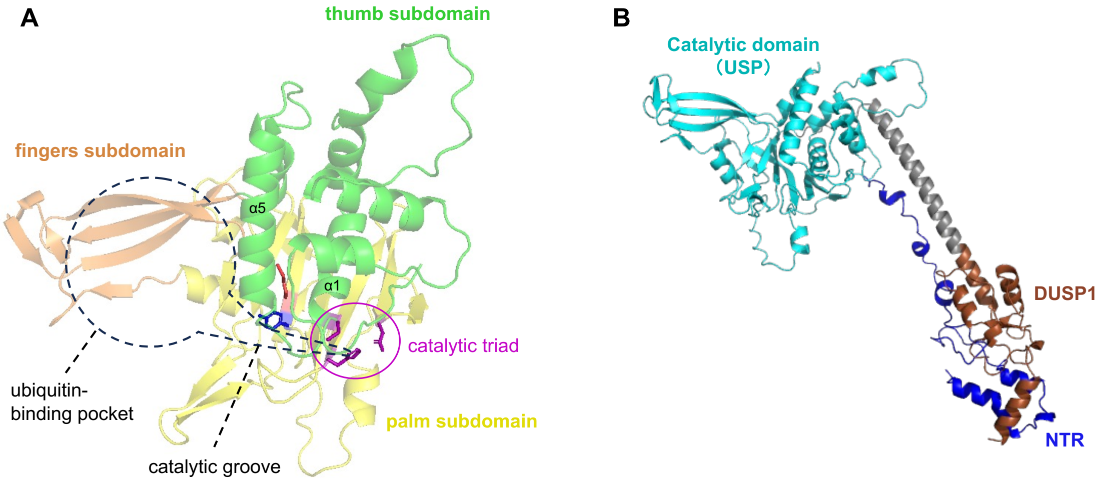


**Fig. S1.** Structure model of USP48.

(A) Predicted structure of the USP domain of USP48 by AF2. The enclosure by black lines indicates the ubiquitin-binding pocket and the catalytic groove. The former captures the ubiquitin core, and the latter interacts with and directs the ubiquitin tail toward the catalytic triad. Magenta, catalytic triad (C98, H353 and N370); blue, Y414; red, M415.

(B) Predicted structure of NTR+USP+DUSP1 (aa 1-568) by AF2. Blue, NTR; cyan, USP domain; brown, DUSP1 domain.

**Supplementary Figure 2**

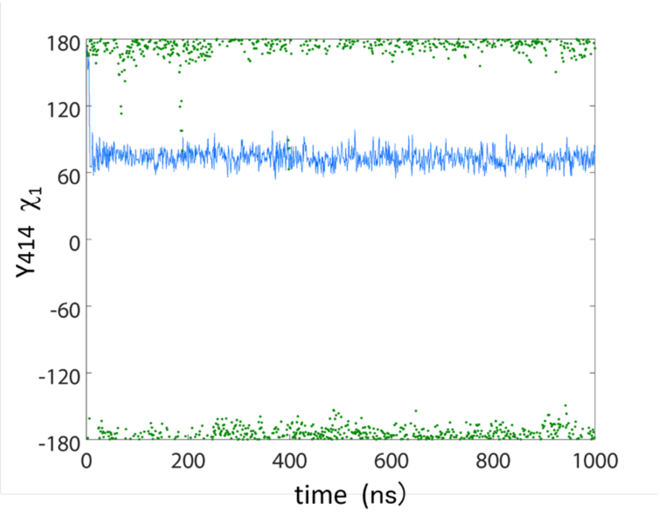


**Fig. S2.** Y414 $\chi_1$ angle as a function of simulation time. USP WT (blue) and USP M415I (green).
